# Multiple allostery in the regulation of PDGFR beta kinase activities

**DOI:** 10.1101/2024.06.06.597829

**Authors:** Yanfeng Zhang, Meimei Wang, Guangcan Shao, Qingbin Shang, Mengqiu Dong, Xiaohong Qin, Li-Zhi Mi

## Abstract

Platelet-derived growth factor receptor beta (PDGFRβ), a type III receptor tyrosine kinase (RTK) with a featured kinase insert, regulates important cellular functions. Dysregulation of PDGFRβ is associated with cardiovascular and fibrosis diseases. Thus, its kinase activity needs to be precisely regulated under physiological conditions. Early studies demonstrated that its kinase was autoinhibited by its juxtamembrane segment and activated by transphosphorylation. However, additional mechanisms are required for the comprehensive regulation of the receptor kinase. Herein, we provide evidences that dimerization of activated kinases, autoinhibition by the kinase insert, and dimerization of inactive kinase, all contribute to the regulation of the receptor kinase. Moreover, we find such multiple allosteric regulation is also conserved in other type III RTKs, including colony stimulating factor 1 receptor (CSF1R). Impairing the allosteric regulation of CSF1R is associated with malfunctions of microglia and demyelination of neurons in Hereditary Diffuse Leukoencephalopathy with Spheroids (HDLS).

## Introduction

PDGFRα and PDGFRβ, in together with CSF-1R, Mast/stem cell growth factor receptor (SCFR or c-Kit), and Fms-like tyrosine kinase 3 (Flt-3) receptors, belong to type III receptor tyrosine kinase subfamily [1–5]. All of them are constituted of 5 extracellular Ig-like domains, a single-pass transmembrane domain (TM), an intracellular juxtamembrane segment (JM), a kinase domain splatted apart by a not-well characterized insert (KI), and a C-terminal tail bearing multiple phosphorylation sites (Fig. 1A) [1–5]. These receptors regulate important biological functions, including cell proliferation, differentiation, and migration. Dysregulation of these receptors is associated with cancer, cardiovascular and fibrosis diseases [1–5]. Thus, a clear understanding of the precise regulation of these receptors is essential.

**Figure 1.**
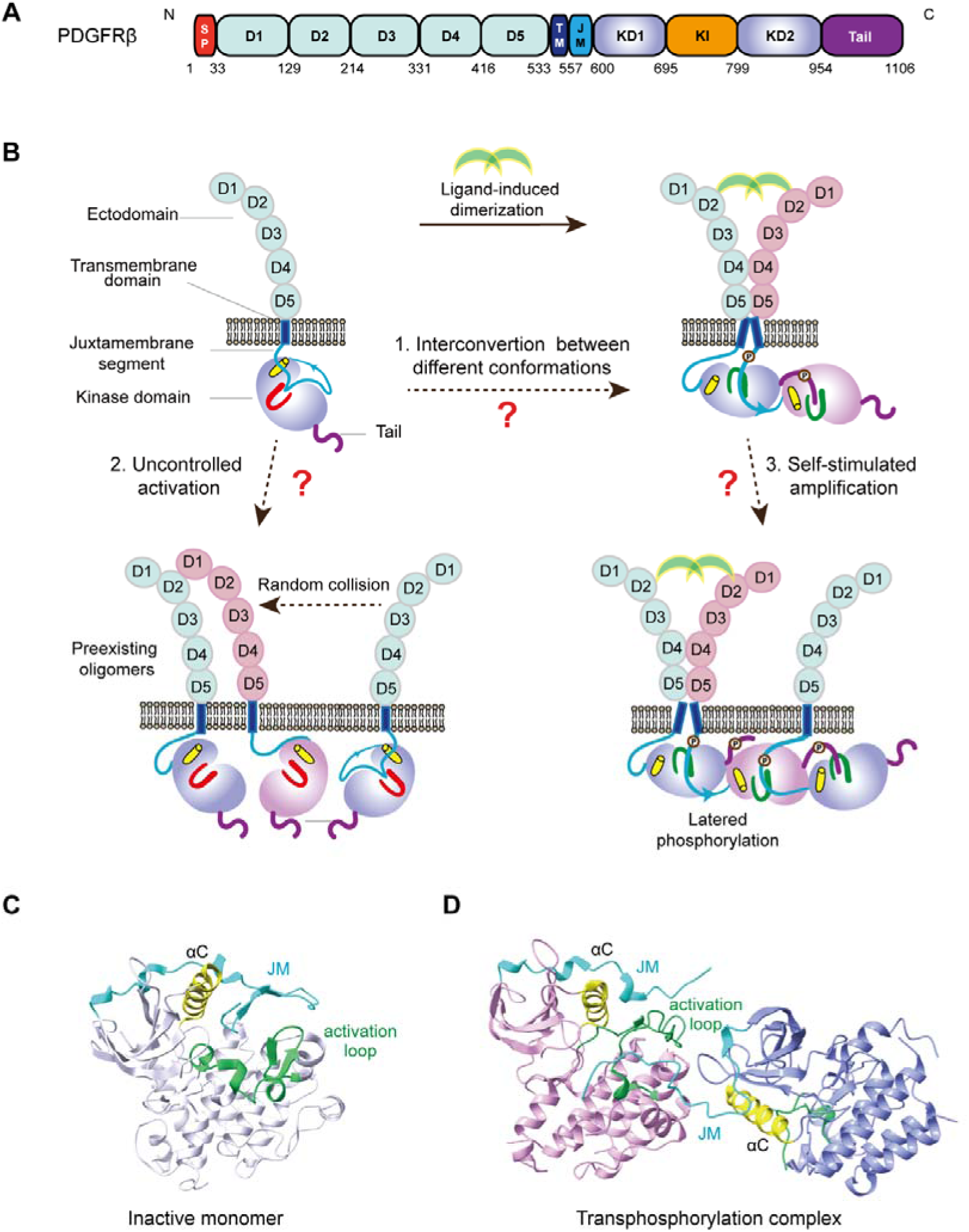
Overview of the structure and activation process of PDGFRβ. (A) Schematic diagram of the primary structure of the PDGFRβ. sp: signal peptide; D1-5: Ig-like ectodomain; TM: transmembrane domain; JM: juxtamembrane segment; KD: Kinase domain; KI: Kinase Insert. (B) Diagram of the activation process of PDGFRβ. Dimerized PDGF ligands bind to two PDGFRβ and activate their kinases. But the precise regulation of the kinases was not fully defined. (C) Autoinhibited state of PDGFRα kinase. Forming an hairpin structure, the JM segment binds to the active site of the kinase and competitively blocks the kinase from binding its substrates. In addition, the JM interacts with the kinase αC helix. Through these interactions, the JM segment stabilizes the kinase in an inactive conformation. (D) Structure of the transphosphorylation complex of c-Kit kinases. The JM of one kinase is attached to the C-lobe of the other and places the tyrosine on the JM into the active site of the latter. When the kinase is activated, such autoinhibition of the JM segment needs to be released.

The receptors in this subfamily were regulated by ligand-induced dimerization and subsequent activation through transphosphorylation (Fig. 1B) [5–8]. In the absence of ligand, these receptors are monomeric and autoinhibited [5, 7–9]. Upon binding to the dimeric ligands, these receptors are assembled into dimers so that two kinases in the dimer could trans-phosphorylate each other and be activated [5, 7, 8].

Crystallographic studies provided further details for these receptor kinases in autoinhibited and transphosphorylation states [10–13]. In the autoinhibited state, the JM of PDGFRα, FLT3, or c-Kit receptor forms an hairpin winding around the αC helix of the kinase and poses the JM tyrosine-containing segment into the active site of the kinase (Fig. 1C) [10, 12, 13]. As such, the JM blocks the kinase from binding to substrate [10, 12, 13]. In the transphosphorylation state of c-Kit kinases, the JM of one kinase is latched onto the C-lobe of another kinase so that the two JM Tyrs from the former kinase can get into the active site of the latter for transphosphorylation (Fig. 1D) [11]. Moreover, in this trans-phosphorylation complex, both kinases are adopted in active conformation [11].

Although these early studies provided a basic framework for understanding the regulation of these receptor kinases, some conceptual gaps regarding the transition and stabilization of different conformational states of the kinase are still missing. For example, it was largely unknown: how is the kinase kept in a precise balance between the autoinhibited state and activated state? In addition, if the kinase is activated merely by auto-phosphorylation, how to prevent uncontrolled trans-phosphorylation and activation which can be introduced constantly by random collision of these receptors on the cell surface? And how to prevent the self-stimulated amplification of external stimuli through lateral phosphorylation of unliganded receptors by ligand-bound receptors (Fig. 1B)?

Accompany with these missing conceptual gaps, mounting experimental evidences also suggested that there were additional layers in the tight regulation of the kinase activities of PDGFRs and their subfamily members. It had been shown that, by rotating the transmembrane helixes of engineered PDGFRβ receptors, the kinases of ligand-bound receptors could be periodically activated [14]. This suggests that bringing the kinases into proximity is not enough to activate the kinases. Moreover, chimeric fusions of PDGFRα or PDGFRβ with proteins in oligomers were found in cancer patients [15–18]. But the kinase activities of these constitutively oligomerized chimeras were not similarly upregulated but were rather depended on the presence of PDGFR JM and other regulatory elements [19–21]. Besides ligand-bound dimers, unliganded PDGFRβ tetramers were also proposed to exist on the cell surface [22, 23]

To find out these missing gaps, we used chemical cross-linking mass spectrometry (CXMS), structure-based mutagenesis, and Rosetta modeling to study the regulation of PDGFRβ kinase activities. We found that the kinase activities of PDGFRβ and its subfamily members were regulated by multiple allostery. These allosteric regulations include: forming an activated symmetric kinase dimer, autoinhibition by the kinase insert, and dimerization of inactive kinases. Disruption of these regulations impaired the functions of these receptors. In addition, we found some HDLS-associated mutations of CSF1R destabilized the dimerization of the active kinases, and thus provided molecular insights for understanding the dominant inheritance of HDLS.

## Results

### A symmetric, active kinase dimer interface was identified in ligand-bound PDGFRβ receptors by CXMS and Rosetta modeling

To find out how PDGFRβ kinases in ligand-bound, dimeric receptors interact with each other to stabilize their activated state, we used CXMS and Rosetta modeling to determine the kinase dimerization interface [24]. The functional full-length PDGFRβ receptor was purified as described [25]. The kinase activities of purified receptors in detergent were analyzed by western-blotting in the presence of none, ATP, PDGF-BB, and Dovitinib. Dovitinib is a PDGFR kinase inhibitor which can stabilize the kinase in the inactive conformation [26, 27]. As it was shown, the kinase activities of purified receptors could be stimulated by the treatment of 0.8 μM PDGF-BB. In addition, the stimulated activities of the receptors could be inhibited by the addition of 10 μM Dovitinib, suggesting purified receptors were functional (Fig. 2A). Then, we cross-linked purified, ligand-bound receptors with 0.8 mM Sulfosuccinimidyl Suberate (BS^3^) or 0.15-0.45 mM Disuccinimidyl Suberate (DSS) for 1 hour. The cross-linked dimers were separated from monomers by SDS-PAGE and analyzed by mass-spectrometry (Fig. 2B).

**Figure 2.**
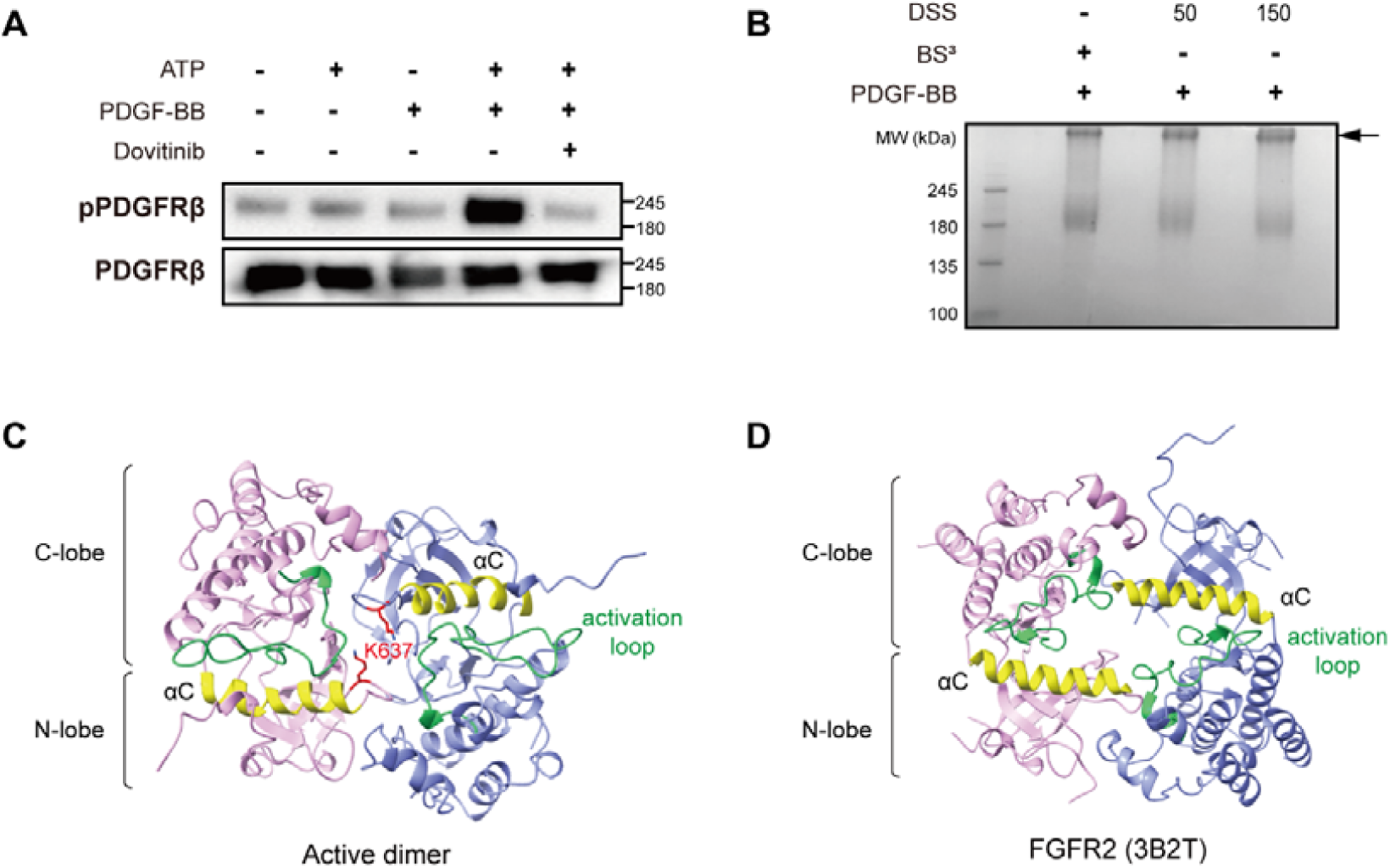
Symmetric dimerization of activated PDGFRβ kinases. (A) Purified PDGFRβ is functional. The activity of purified PDGFRβ was analyzed in the presence and absence of ATP, PDGF-BB, and Dovitinib. The phosphorylation and protein levels of the samples were analyzed by western blotting using 4G10 and protein C antibodies, respectively. (B) Cross-linking of purified PDGFRβ with BS^3^ and DSS. The purified PDGFRβ was treated with PDGF-BB at 4 °C for 1.5 h, and then the mixture was cross-linked with BS^3^ (1 μg BS^3^/μg protein) or DSS (The molar ratio of protein to DSS is 1:50 or 1:150) at room temperature for 1 h. The reaction was stopped with 20 mM NH_4_HCO_3_. The samples were separated by SDS-PAGE. The arrow indicates cross-linked bands of PDGFRβ. (C) Dimeric structures of activated PDGFRβ kinases. The model of activated PDGFRβ kinase dimer was calculated with Rosetta. K637s were shown in red sticks. The αC helix is in yellow; and the activation loop is in green. (D) The crystallographic dimer of FGFR2 kinase in active conformation (PDB 3B2T). (E) Sequence alignment of PDGFRβ, CSF1R and FGFR2. Residues at the kinase dimerization interface of PDGFRβ and FGFR2 were marked with green squares and purple diamonds, respectively. Residues at CSF1R kinase dimerization interface were marked with yellow and red stars with red stars emphasizing the mutations associated with HDLS.

Nine cross-linked peptide pairs were identified from ligand-bound dimeric PDGFRβ receptors (Table 1, Fig. S1). Especially, two Lys387s on the extracellular subdomain D4s were cross-linked together in ligand-bound receptors (Table 1). This is consistent with the experimental evidence that the proximity of D4 is critical for PDGFRβ activation. In addition, it is compatible with the crystal structure of ligated c-Kit ectodomain (PDB 2E9W) [28, 29], suggesting our cross-linking experiments are valid.

**Table 1.** Cross-linked peptide pairs of ligand-bound PDGFRβ.

| PDGFRβ | | PDGFRβ | | Spectral counts | Best E value |
| --- | --- | --- | --- | --- | --- |
| Sequence | Cross-linked lysine | Sequence | Cross-linked lysine |  |  |
| VTDPQLVVTLHEKK | 163 | SYICKTTIGDR | 191 | 2 | 4.49E-04 |
| • VKVAEAGHYTMR | 387 | VKVAEAGHYTMR | 387 | 8 | 4.83E-04 |
| MLKSTAR | 637 | MLKSTAR | 637 | 3 | 8.32E-05 |
| • MLKSTAR | 637 | SSEKQALMSELK | 645 | 4 | 5.36E-04 |
| • MLKSTAR | 637 | DSNYISKGSTFLPLK | 860 | 1 | 3.39E-07 |
| MLKSTAR | 637 | KYQQVDEEFLR | 969 | 3 | 3.78E-05 |
| VAVKMLK | 634 | SSEKQALMSELK | 645 | 4 | 9.07E-04 |
| SSEKQALMSELK | 645 | DSNYISKGSTFLPLK | 860 | 1 | 4.13E-03 |
| • LLGEGYKK | 967 | KYQQVDEEFLR | 969 | 3 | 4.09E-05 |

At intracellular side, we reasoned that the K637-K637, K637-K969, and K967-K969 represent the inter-molecularly cross-linked pairs. In the threading model of PDGFRβ kinase, which is based on the structure of active c-Kit kinase (PDB 1PKG), K969 is located too far away from K637 to be intra-molecularly cross-linked. Similarly, K969 is located too close to K967 to be intra-molecularly cross-linked. Other cross-linked Lys pairs were individually screened *in silico* as inter- or intra-molecular pairs according to the criteria described in the method section.

By using the proximities of inter-molecularly cross-linked Lys pairs as distance constrains, we calculated a model of activated PDGFRβ kinase dimer using Rosetta Docking protocols [24]. As K637-K637, K637-K969, and K967-K969 were reasoned to be inter-molecularly cross-linked pairs, they impose a nearly 2-fold symmetry constrain on the dimeric kinase assembly. As such, we used a 2-fold symmetry constrain in the global docking step and released such constrain in the final local-docking step.

The calculated active PDGFRβ kinase dimer was symmetric (Fig. 2C). The total buried surface area of the dimer was 2290 Å^2^. The binding energy and the P-value of the assembly was estimated to be −9.0 kcal/mol, and 0.315, respectively, using the PISA server [30]. In this dimer, two kinases were interlocked together through the interactions between the N-lobe residues (G603, R604, T605, S608, Q613, S638, T639, A640, S642, K645, G674 and G675) and C-lobe residues (R830, P866, L867, K868, W869, P905, E906, P908 and M909).

This symmetric dimer is reminiscent of the crystallographic dimer of FGFR2 kinase (PDB 3B2T) (Fig. 2D), in which the activity-compromised mutant of FGFR2 kinase adopts an active conformation [31]. Similar symmetric assembly was also found in the crystal structures of the active RET (PDB 2X2L) and FGFR1 kinases (PDB 5FLF) (Fig. S2) [32, 33]. Moreover, the dimer interface was well-conserved among the type III receptor tyrosine kinases [34] (Fig. S3).

### Disruption of the symmetric kinase dimer interface abolished the ligand-stimulated activities of PDGFRβ and CSF1R

To validate the physiological relevance of identified symmetric kinase dimer interface, 11 out of 21 residues buried at the PDGFRβ kinase dimer interface were randomly selected and mutated (Fig. 3A). Except for P908A, all other mutations impaired the stimulated activity of PDGFRβ (Fig. 3B). The R604E, G603E, A640E, and K645E mutations exhibited moderate effects, while other mutations had more significant impact on activity. The residues R830 and P866 were near the ATP binding pocket, whereas the other residues were not involved in ATP or substrate binding.

**Figure 3.**
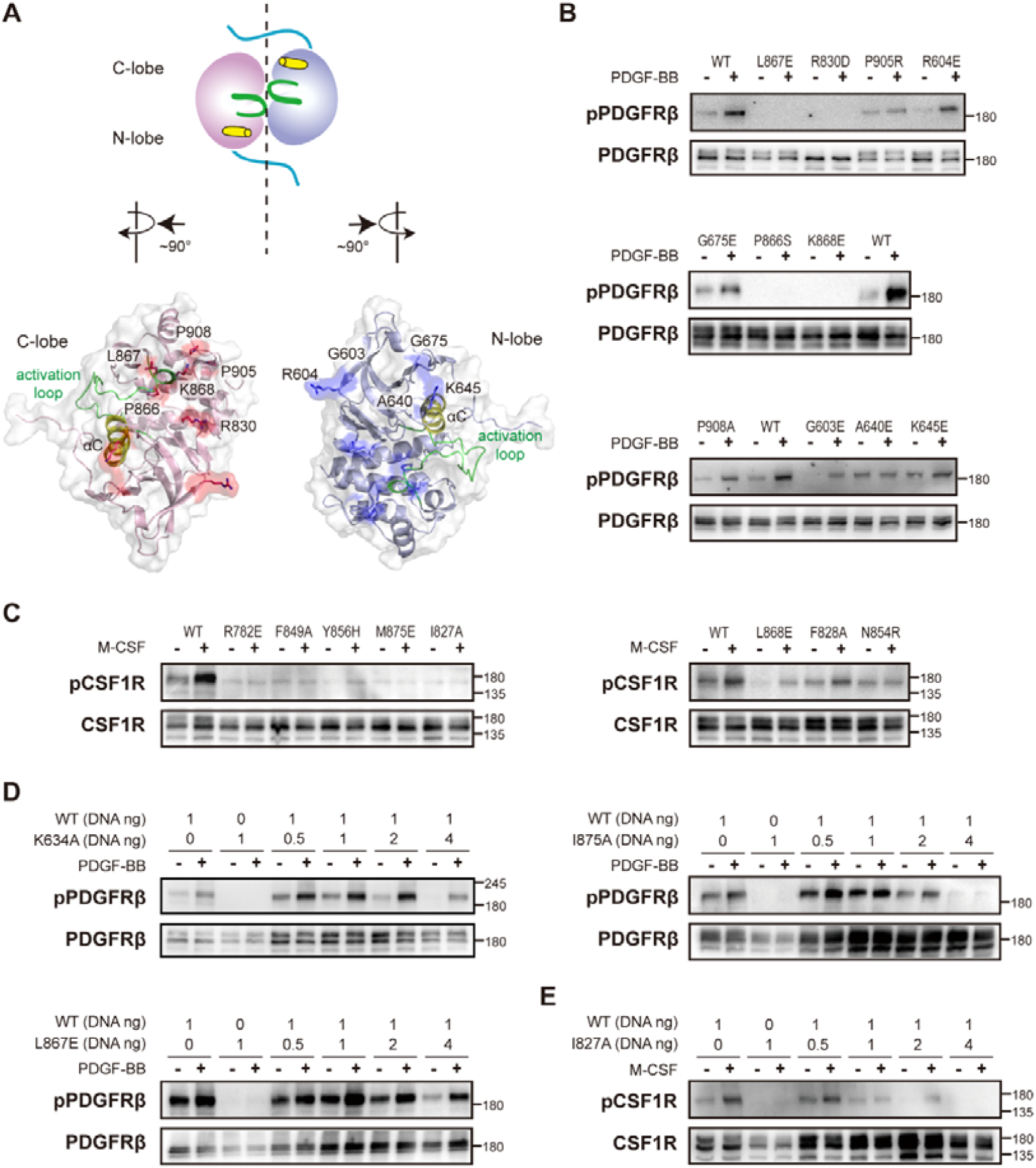
Disruption of activated kinase dimerization impairs ligand-stimulated activation of PDGFRβ. (A) Calculated structural model for activated PDGFRβ kinase dimer. For clarity, two activated kinases in the dimer were rotated clockwise and counterclockwise by 90°, respectively. N-lobe residues at the dimer interface were shown in blue sticks on the right, while C-lobe residues at the dimer interface were shown in red sticks on the left. (B) Mutations of the residues at the symmetric dimer interface impair the activation of PDGFRβ. Eleven residues at the symmetric kinase dimer interface were selected and mutated. Except for P908A, every mutation reduced the stimulated activity of PDGFRβ. (C) Mutations on CSF1R kinase dimer interface are linked to the pathogenesis of HDLS and impaired kinase activation. All selected mutations impair the ligand-dependent activation of CSF1R. (D) The impact of the expression levels of PDGFRβ mutants on the activity of co-expressed WT receptor. The inactive mutants of PDGFRβ were co-transfected with the WT receptor. In the transfection, the amount of the plasmid encoding WT PDGFRβ was fixed, while the amount of the plasmid encoding the inactive mutant was increased from 0.5 to 4 folds of the WT. Increasing the expression of I875A mutant could proportionally inhibit the stimulated activity of WT receptor. (E) The impact of the expression levels of CSF1R mutant on the activity of co-expressed WT receptor. The inactive mutant of CSF1R, I827A, was co-transfected with the WT receptor. In the transfection, the amount of the plasmid encoding WT I827A was fixed, while the amount of the plasmid encoding the I827A was increased from 0.5 to 4 folds of the WT. Increasing the expression of I827A could proportionally inhibit the stimulated activity of WT receptor.

Strikingly, the conserved kinase dimer interface was overlapping with CSF1R mutations associated with HDLS, a rare autosomal dominant neurodegenerative disease [35–37]. We hypothesized that these disease-associated mutations would affect the stability of activated kinase dimer, and thus would impair the stimulated activity of CSF1R. To test this hypothesis, eight residues associated with HDLS were mutated and all of them impaired the stimulated activity of CSF1R (Fig. 3C).

It is unlikely that our selected mutations all interfere with substrate recognition in trans-phosphorylation. But we carefully examined such possibility by co-transfecting each selected mutant with the wild-type receptor. In co-transfection experiments, the K634A/WT combination was used as a control. K634 is the key residue required for catalyzing the phospho-transfer reaction. Presumably, K634A mutation should have little effect on the equilibrium of the kinase conformations. If auto-phosphorylation and dimerization of activated kinases were all required for PDGFRβ activation, co-expression of WT receptor with a mutant that can impair both auto-phosphorylation and active kinase dimerization should have a stronger impact on kinase activity than co-expression of K634A mutant with WT receptor. Indeed, we found the stimulated activity of co-expressed WT and I875A was lower than that of co-expressed WT and K634A (Fig. S4). Our structural analysis revealed that I875 residue is located on αEF helix, in proximity to dimerization interface. In addition, I875 is predicted to be at the transphosphorylation interface.

To further validate this result, we did a titration experiment. In the transfection, we fixed the amounts of the plasmids encoding the WT PDGFRβ but increased the amounts of the plasmids encoding the mutants (Fig. 3D). With the increasing amounts of the plasmids encoding I875A mutant, the stimulated activities of the co-transfected I875A/WT receptors decreased proportionally. In comparison, the stimulated activities of co-transfected L867E/WT as well as K634A/WT receptors were not decreased proportionally.

The above experimental evidences are consistent with our calculated structure of activated PDGFRβ kinase dimer. Based on our structure, I875 is located at the substrate recognition site of the kinase. Moreover, I875A mutation could destabilize the active kinase dimerization. As such, I875A mutation generates a stronger impact on the kinase activity of co-transfected receptors than K634A mutation.

I875 is conserved across the type III RTKs (Fig. S3). Mutation of its equivalent residue in CSF1R, the I827, is linked to the onset of HDLS [37]. To investigate the significance of this mutation on the pathogenesis of HDLS, we mutated CSF1R I827 to Ala and performed the titration experiment as described above (Fig. 3E). The stimulated activities of the samples co-transfected with I827A and WT CSF1Rs were reduced depending on the amount of co-transfected I827A plasmids. This result indicated that I827A mutation could inhibit the activity of WT CSF1R in *trans*, which had implications for dominant heritance of this disease.

### The kinase insert could autoinhibit the ligand-induced activation of PDGFRβ

To understand how inactive kinases are associated together in ligand-bound PDGFRβ receptors, we used a PDGFRβ kinase inhibitor, Dovitinib, to stabilize the kinase in the inactive conformation (Fig. 4A, 4B) and studied the kinase association by CXMS [38]. Purified PDGFRβ receptors treated with both PDGF-BB and Dovitinib showed a different cross-linking spectrum from the sample treated with PDGF-BB alone (Table 2, Fig. S5). Especially, many cross-linking pairs were identified between the kinase insert and the kinase domain. In addition, the cross-linking pair between the two K387s on the receptor extracellular D4 domains was kept, indicating the proximity of the D4 domains in ligand-bound PDGFRβs was not affected by stabilizing the kinase in an inactive conformation.

**Figure 4.**
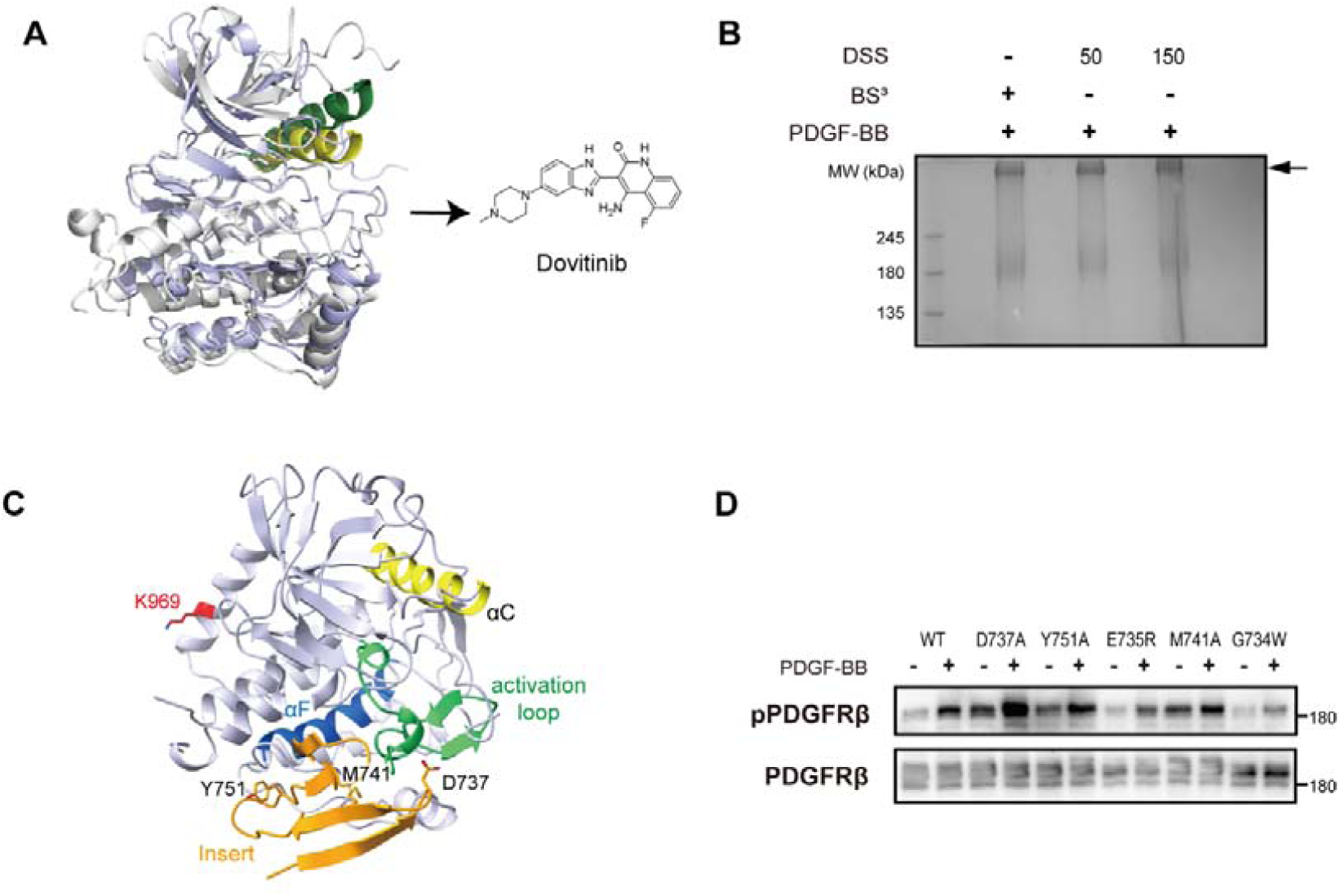
Autoinhibition of PDGFRβ by its kinase insert. (A) PDGFRβ kinase can be stabilized in an inactive conformation by Dovitinib. Superimposition of the structure of activated PDGFRβ kinase with that of inactive FGF1 kinase complexed with Dovitinib (PDB 5A46). The activated PDGFRβ is in light blue, and its αC helix is in yellow; the inactive FGF1 is in gray, and its αC is in green. (B) Dovitinib-bound PDGFRβ was cross-linked by BS^3^ and DSS. Purified PDGFRβ was incubated with PDGF-BB at 4 °C for 1.5 h, and then was treated with 10 μM Dovitinib for 5 min. The mixtures were crosslinked with BS^3^ or DSS at room temperature for 1 h. The arrow indicates cross-linked bands of PDGFRβ. (C) Schematic diagram of PDGFRβ kinase insert. Using cross-linking data as constraints, the structure of the kinase insert was docked onto the inactive kinase structure using Rosetta. The kinase insert was docked to the kinase C-lobe nearing the αF helix. The binding site of the kinase insert overlaps with the substrate recognition site and the activated kinase dimer interface. (D) Mutations on the kinase insert could impair the activity of PDGFRβ. Five KI residues interacting with the kinase were mutated. Three of them (D737A, Y751A, and M741A) could enhance the activity of the kinase.

**Table 2.**
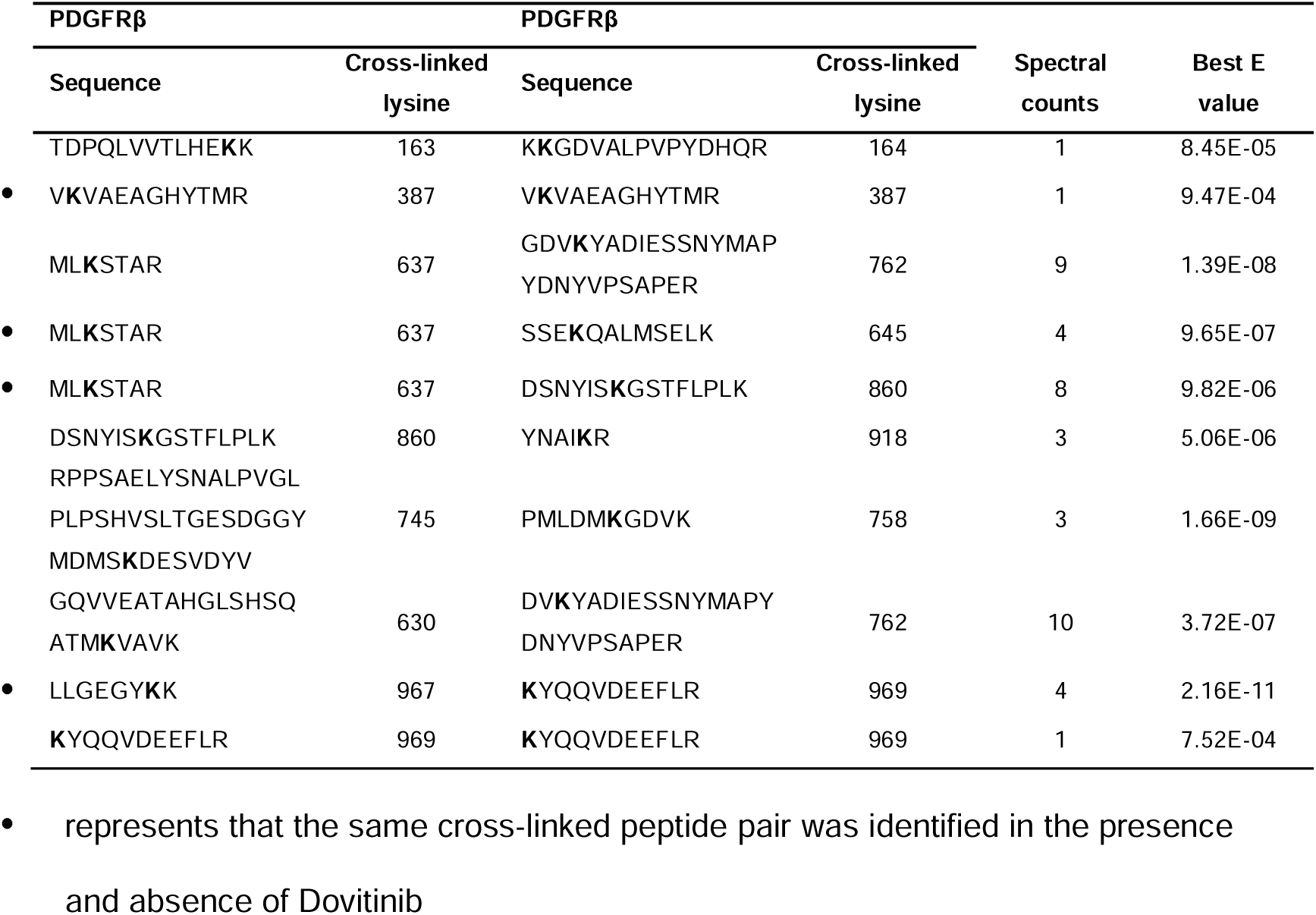
Cross-linked peptide pairs of ligand-bound PDGFRβ inhibited by Dovitinib.

To model the inactive kinase association in ligand-bound, Dovitinib-inhibited receptors, we took three steps in Rosetta computation: *Ab initio* modeling of the kinase insert, docking of the insert core onto the kinase, and docking the insert-bound inactive kinases [7].

In the *Ab initio* modeling of the kinase insert, all-atom models were generated and refined [24]. These models varied significantly in their structures. In the top cluster, the RMSDs of calculated models were converged to 8 Å (Fig. S6A). However, these models shared a conserved core structure composed of 4 β-strands. The RMSDs of the core structures were converged to 1 Å (Fig. S6B). In addition, the sequence of this kinase insert core is well-conserved across Metazoa (Fig. S7).

Using the cross-linking data and the calculated structure of the insert core, we modeled the association between the insert and the inactive kinase domain with Rosetta [5]. In docking models, the insert core was mapped to the kinase C-lobe nearing the αF helix (Fig. 4C). In this position, the kinase insert could sterically block the kinase from recognizing its substrates and forming the activated kinase dimer.

To verify this point, we mutated 5 residues on the kinase insert, which interact with the kinase domain in our model (Fig. 4D). In comparison with the WT receptor, the stimulated activities of 3 out of 5 mutants were significantly enhanced, while the activities of the others remained similarly, confirming the autoinhibitory function of the kinase insert (Fig. 4D).

Unfortunately, we could not unambiguously determine the interface for inactive kinase dimerization due to the limited number of identified cross-linking pairs (Table 2). However, two Lys969s on the kinase C-terminal αJ helices were cross-linked, suggesting that these two helices should be in proximity to each other in the inactive kinase dimer. This result is consistent with the report that the C-terminus of PDGFRβ is only exposed and detectable by a conformational-specific antiserum in the presence of ligand [39].

### Formation of a tetrameric complex is not required for the transactivation of PDGFRβ kinase

As it had been proposed that PDGFRβ could exist as a tetramer on the cell surface, we examined such possibility by analyzing the crystal packing of known RTK III kinase structures [22, 23]. From these structures, we found activated c-Kit kinases could form a circular tetrameric complex in transphosphorylation through four nearly identical head-to-tail interactions (PDB 1PKG) (Fig. 5A) [11]. The interface of these interactions is well-conserved among RTK III kinases [34] (Fig. 5B, S3), indicating this assembly might be functionally relevant.

**Figure 5.**
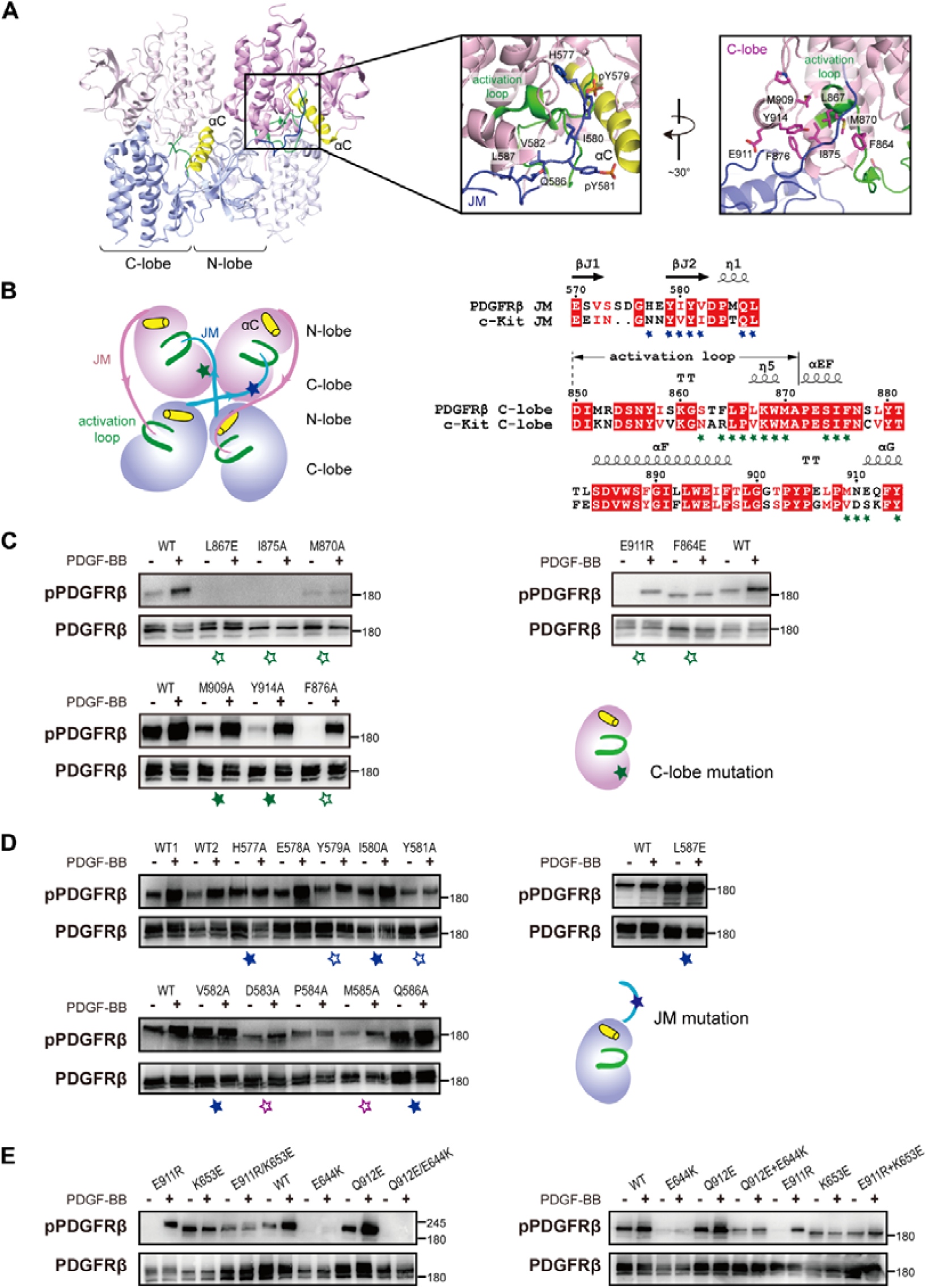
Tetrameric assembly of PDGFRβ is not required for its ligand-dependent activation. (A) The crystal packing of tetrameric c-Kit kinases in transphosphorylation. Residues at the tetrameric interface were shown in close-up views. JM residues were shown in blue sticks; and C-lobe residues were shown in purple sticks. (B) Model of tetrameric PDGFRβ kinases and the sequence alignment of c-Kit and PDGFRβ. The tetrameric interfaces are located at two different regions: the JM segment and the C-lobe. Residues at the JM interface were shown in blue stars; and residues at the C-lobe interface were shown in green stars. (C) The effect of mutations at the C-lobe interface on PDGFRβ activation. Mutants with decreased activity were marked with hollow green stars. (D) The effect of mutations at the JM interface on PDGFRβ activation. JM mutants with decreased activity were marked with hollow blue stars. Residues, which are not at the interface but have decreased activity, were marked by hollow purple stars. (E) Double mutations of electrostatic interaction pair could not restore the impaired kinase activity of singly-mutated receptors. Residues forming electrostatic interactions at the tetrameric interface were mutated to the residues with opposite charges. A pair of non-interacting residues (644/912) was used as a control. Left, comparison of the activities of single mutants with double mutants. Right, comparison of the activities of single mutants with those of paired co-transfected single mutants.

So, we tested whether the formation of this tetramer was required for stabilizing PDGFRβ kinases into activated dimer-to-dimer interactions in transactivation. To this end, we mutated a series of residues at the transphosphorylation interface. Based on the localization of these residues, they could be divided into two different groups: one is at the kinase C-lobe, and the other is at the juxtamembrane (Fig. 5B). Six out of eight mutations at the kinase C-lobe dramatically reduced ligand-stimulated activity of PDGFRβ (Fig. 5C). In contrast, only 2 (Y579A and Y581A) out of 7 JM mutations at the interface reduced ligand-stimulated activity of PDGFRβ (Fig. 5D), while three JM mutations (V582A, Q586A and L587E) at the interface enhanced the kinase activity. We also found that 2 other mutations (D583A and M585A) not at the interface could reduce the kinase activity (Fig. 5D). These results in together do not support the possibility that forming this transphosphorylation tetramer is required for PDGFRβ transactivation.

To further validate this conclusion, we introduced an electrostatic interaction pair (911/653) into the transphosphorylation interface and studied whether double mutations could restore the impaired activities of singly-mutated receptors. As a control, a pair of non-interacting residues (644/912) was selected and mutated. Except for Q912E, every single mutation reduces the stimulated activity of the kinase. But none of the double mutations can restore the kinase activity to the averaged level of the two singly-mutated receptors (Fig. 5E, left). Similar results were obtained when we compared the averaged activity of two single mutants with that of paired co-transfected single mutants (Fig. 5E, right).

These results suggested that forming the tetrameric transphosphorylation complex is not required for the transactivation of PDGFRβ.

## Discussion

The regulation of PDGFRβ kinase activity was far more complicated than what we had expected. Besides previously identified autoinhibition by JM and activation by transphosphorylation [5, 6, 12], we found additional regulatory layers that are required for the precise regulation of the activities of PDGFRβ and its subfamily members. These regulatory layers include: formation of a specific activated kinase dimer, autoinhibition by the kinase insert, and formation of an inactive kinase dimer (Fig. 6). Identification of these additional layers would clarify the comprehensive regulation of PDGFRβ and other receptors in its subfamily.

**Figure 6.**
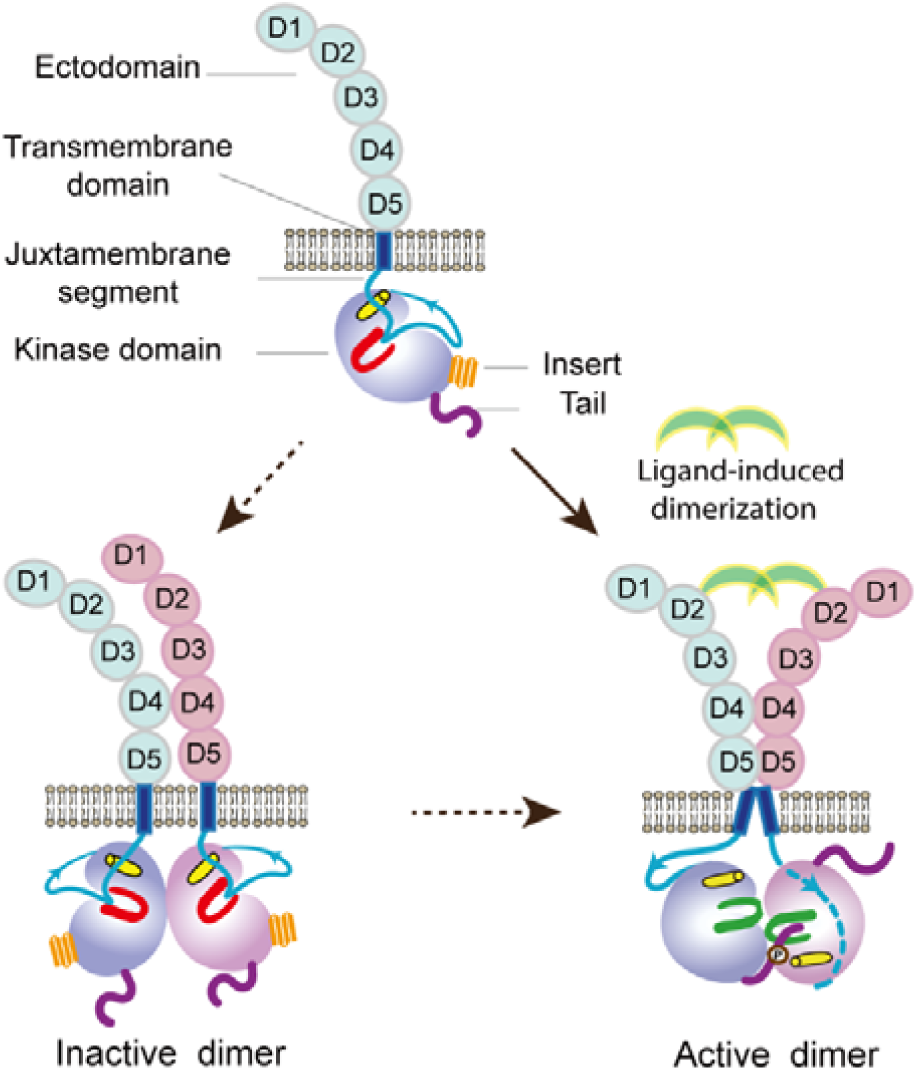
Schematic diagram for a potential mechanism of PDGFRβ activity regulation.

Allosteric activation might be a general theme in the regulation of receptor tyrosine kinases. Such mechanism had been demonstrated in the study of EGFR, a non-canonical receptor tyrosine kinase, by Kuriyan *et. al.* [40, 41]. Unlike PDGFRβ, phosphorylation of EGFR kinase activation loop is not required for its activation.

Instead, two kinases of ligated EGFRs form a cyclin/CDK-like asymmetric dimer to activate the “CDK-like” receiver kinase [40, 41]. A similar asymmetric dimerization was also found in the tetrameric complex of c-Kit kinases [11], but such asymmetric interaction was not required for the activation of PDGFRβ. Instead, two activated kinases in ligand-bound PDGFRβ receptors form a symmetric dimer. This symmetric dimer interface is well-conserved among the type III RTKs, including CSF1R (Fig. S3). Similar symmetric dimerization could be found in the crystal packing of FGFR1, RET, and FGFR2 kinases [32, 33, 42].

Asymmetric and symmetric dimerization have different impacts on receptor oligomerization and signaling. Two distinct interfaces of asymmetric dimerization allow the receptors to be assembled continuously, forming a linear polymer for lateral signal propagation [40, 41]. But if these two distinct interfaces were used for symmetric dimerization, receptors could only be assembled into dimers but not high-order oligomers. This self-constrained dimerization could prevent uncontrolled signaling caused by random collision of receptors, which happens constantly on the cell surface. In addition, it could limit the signal self-amplification through lateral phosphorylation. Indeed, many kinases, including Ser/Thr kinases, are assembled into symmetric dimers in their signaling [43].

Dysregulation of receptor tyrosine kinases is often associated with various human diseases, including cancer and neurodegenerative disease [1, 3]. While many cancers are associated with gain-of-functional mutations of kinases, neurodegenerative diseases are often associated with loss-of-functional mutations of kinases [35, 36]. HDLS, a dominant neurodegenerative disorder, is caused by microglia dysfunction, which in turn leads to demyelination of neurons [36]. Many missense mutations of CSF1R had been identified in HDLS patients [36]. But their impacts on CSF1R functions and their underlying pathogenic basis were largely unknown [35, 36]. In our studies, we found that some HDLS-associated CSF1R mutations impaired the stimulated activity of the receptor and the dimerization of the activated kinases. In addition, I827A mutation does not only impair the kinase activity of itself own but could also inhibit the activity of co-expressed WT receptor in a dose dependent manner. Our results provide molecular insights for understanding the regulation of CSF1R function in microglia and the molecular basis for the dominant heritance of HDLS.

A striking feature of the type III RTKs is the presence of a kinase insert that splits the kinase domain into two separated halves [3, 5]. The KI of PDGFRβ had been shown to be important for PDGF-induced mitogenesis [44, 45]. However, less was known about the structures and functions of these KIs in kinase activity regulation. In limited studies on the KIs of type III RTKs, controversial results had been reported [44, 46]. To investigate the biological functions of PDGFRβ KI, we used Dovitinib to stabilize the PDGFRβ kinase in an inactive conformation. We found the KI of PDGFRβ could sterically block substrate recognition and dimerization of activated kinases. As such, the KI of PDGFRβ autoinhibits the activation of the receptor. However, our studies were limited by the uncertainty of the complete structure of the KI. Further studies are required to define the atomic details of the KI and its interactions with the splatted kinase domain.

Another limitation of our study is that the inactive kinase dimer interface was not defined. However, a clear contrast between Dovitinib-bound and unbound kinase dimer interactions was detected in CXMS experiments. From limited cross-linking pairs detected in the inactive kinase dimer, we found the two C-terminal αJ helices should be in proximity to each other.

In summary, we defined that multiple allostery is involved in the precise regulation of the activities of PDGFRβ and its subfamily members. Disruption of these regulatory layers could impair the functions of these receptors, which might have clinical implications.

## Experimental procedures

### Materials

Antibodies were obtained from the following sources: Protein C-Tag Antibody (HPC4) (Rabbit polyclonal) was from Genscript (Piscataway, NJ, USA); Anti-Phosphotyrosine Antibody, clone 4G10 (Mouse Monoclonal) was from Merck Millipore (Darmstadt, GER); Human PDGF-BB (PDB-H4112) and Human M-CSF/CSF-1 (MCF-H5218) were from ACRO Biosystems (Newark, DE, USA); Dovitinib was from MedChemExpress (Monmouth Junction, NJ, USA); Disuccinimidyl Suberate (DSS) and Sulfosuccinimidyl Suberate (BS^3^) were from Thermo Fisher Scientific (Waltham, MA).

### Construction of plasmids

The genes encoding wild-type full-length human PDGFRβ (from Dr. Jiahuai Han laboratory) and CSF1R (from Sino Biological Inc) were subcloned into the pEF1-puro plasmid (from Dr. Timothy A. Springer) at *Bam*HI and *Xho*I restriction sites. All mutations were constructed by using Gibson assembly kit (NEB). The signal peptide of murine immunoglobulin κ chain was used for expression of PDGFRβ and CSF-1R receptors. For purification and western blotting, a protein C and a SBP-tag were fused tandemly to the C-terminus of PDGFRβ or CSF-1R receptor. For monitoring the transfection efficiency, an IRES element and EGFP gene were added after the receptor coding sequence. All constructs were verified by Sanger sequencing.

### Cell culture, transfection and Immunoblotting

HEK293T cells were cultured in Dulbecco’s modified Eagle’s medium (DMEM) supplemented with 10% FBS at 37 °C with 5% CO_2_. When cells reached 40-50% confluency, 0.5 μg of plasmid was mixed with 1.5 μg linear polyethyleneimine (PEI), and then added into a well of cultured HEK293T cells in 12-well plates. After transfection of the cells for 5 hours, the culture media was replaced with serum free DMEM. After 20 hours of post-transfection, cells were treated with 50 ng/μL PDGF-BB or CSF (from ACRO Biosystems) for 20 minutes in CO_2_ incubator. The treated cells were lysed and prepared for western blotting analysis as described [25]. The expression of PDGFRβ and CSF1R was detected by western-blotting using anti-protein C antibody (Genscript, USA). The phosphorylation levels of PDGFRβ or CSF1R were detected by western-blotting using 4G10 antibody (Merck Millipore, Germany).

### Expression, purification of PDGFRβ and Kinase assay in vitro

To express PDGFRβ, HEK293F cells at 1.5×10^6^ cells/mL were seeded into 1 L SMM-293TI medium in a 2 L conical flask rotating at 120 rpm/min in a 5% CO_2_ incubator at 37 °C. One mg plasmids and 3 mg PEI were mixed and incubated at room temperature for 30 min. Then, the mixture was added into the flask. After 3 days, the transfected cells were harvested and stored at −80 °C.

To purify PDGFRβ, harvested cells were resuspended and lysed as described [25]. The lysate was cleared by ultracentrifugation at 50,000 ×*g* for 30 min at 4 °C. The cleared lysate was loaded onto a strep-tactin column (IBA, Germany) and purified as described [25]. In chemical cross-linking experiments, the lysis buffer was replaced by 50 mM HEPES, pH 8.0, 400 mM NaCl, 1 mM EDTA, 10% glycerol, 0.15% Triton X-100, 100 mM Na_3_VO_4_, 0.5 mM TCEP, and 1× Complete protease inhibitor cocktail. The wash buffer was replaced by 50 mM HEPES, 150 mM NaCl, 1 mM EDTA, 10% glycerol, 0.1% Triton X-100, 0.5 mM TCEP, and 2 mM PMSF.

In kinase activity assay, purified PDGFRβ was treated with 0.8 μM PDGF-BB in 25 µL of reaction buffer containing 8 mM HEPES and 160 µM ATP on ice for 30 min. The reaction was stopped by adding 6× SDS sample buffer. To inhibit PDGFRβ kinase activity, 10 µM Dovitinib in together with 0.8 μM PDGF-BB was added to the purified receptor on ice for 5 min. The expression and phosphorylation levels of PDGFRβ were detected by western blotting using anti-protein C and 4G10 antibodies, respectively.

### Chemical Cross-linking of proteins coupled with mass spectrometry (CXMS) analysis

The purified protein was concentrated by using an Amicon Ultra 0.5 mL 50 kDa filter (Millipore) at 12,000 ×*g* for 10 min. The protein concentration was calibrated by using BSA as standard (∼ 0.3-1 mg/mL).

About 8 μg of purified PDGFRβ was incubated with 0.18 μM PDGF-BB at 4 °C for 1.5 h. Then, the proteins were crosslinked with BS^3^ (1 μg BS^3^ per 1 μg protein) or DSS (the molar ratios of protein to DSS are 1:50 and 1:150) for 1 h at room temperature. The reaction was quenched by adding 20 mM NH_4_HCO_3_. For stabilizing the kinase of PDGFRβ in an inactive conformation, 10 μM Dovitinib was added to the protein solution on ice for 5 min. The cross-linking experiment was performed similarly as described.

Next, the cross-linked proteins were precipitated with 4 volumes of ice-cold acetone, resuspended in 8 M urea, 100 mM Tris pH 8.5, and digested with 0.16 μg Trypsin (Promega) in 2 M urea, 100 mM Tris (pH 8.5). The LC-MS/MS analysis was performed on an Easy-nLC 1000 II HPLC (Thermo Fisher Scientific) coupled to a Q-Exactive HF mass spectrometer (Thermo Fisher Scientific). Peptides were loaded and desalted on a pre-column (75 μm ID, 6 cm long, packed with ODS-AQ 120 Å–10 μm beads from YMC Co., Ltd.), and subsequently separated on an analytical column (75 μm ID, 12 cm long, packed with Luna C18 1.9 μm 100 Å resin from Welch Materials). The sample on the analytical column was eluted with a linear reverse-phase gradient from 100% buffer A (0.1% formic acid in H_2_O) to 30% buffer B (0.1% formic acid in acetonitrile) over 56 min at a flow rate of 200 nL/min. The top 15 most intense precursor ions from each full scan (resolution 60,000) were isolated for HCD MS2 (resolution 15,000; normalized collision energy 27) with a dynamic exclusion time of 30 s. Precursors with 1+, 2+, 7+ or above, or unassigned charge states were excluded. The pLink 2 [47] software was used to identify cross-linked peptides with precursor mass accuracy of 20 ppm and fragment ion mass accuracy of 20 ppm. The results were filtered by applying a 5% FDR cutoff at the spectral level and then an E-value cutoff at 0.001 [48].

### Structural Modeling

The global docking of the PDGFRβ kinase dimer was calculated with Cluspro [49]. The local docking of the kinase dimer was calculated with Rosetta as described [50]. Each cross-linked residual pair was used individually as a distance constrain in the calculation of kinase dimer. In the calculation, the pair-wise distance was calculated with Xwalk [51]. 500 models with the lowest energy scores were selected. Then, these models were screened for their satisfactions of cross-linking constrains (< 30 Å) and the requirement for buried interface area (> 900 Å^2^). Two largest clusters were selected from these screened models. The structural models calculated with each distance restrain were compared. Only those pairs that generate similar, converged structural models were selected as inter-molecular constrains for next round calculation. Next, every two selected cross-linked pairs were combined as two distance constrains in the local docking calculation. Those that gave converged models were selected. Finally, three out of five cross-linked pairs were combined as inter-molecular distance constrains in the calculation of the final docking models.

## Data availability

Further information and requests for resources and reagents should be directed to and will be fulfilled by the lead contact, Li-Zhi Mi Original images of SDS-PAGE gels and WBs reported in this paper will be shared by the lead contact upon request.

## Supporting information

This article contains supporting information.

## Acknowledgments

The authors are grateful to Dr. Jiahuai Han (Xiamen University) for providing the cDNA of human PDGFRβ.

## Funding and additional information

LZM was funded by National Natural Science Foundation of China (#31670738, #31470730). XQ was funded by Tianjin Municipal Science and Technology Bureau and Tianjin Natural Science Foundation (# 21JCQNJC01660).

## Conflict of interest

The authors declare no conflicts of interest.

## Notes

### Competing Interest Statement

The authors have declared no competing interest.

### Summary of Updates

Revise page 9, lines 4-5; Funding information updated.

