## Supplemental Figures S1-S7 for "Multiple allostery in the regulation of PDGFR beta kinase activities"

**Contents**:

Supplemental Figures S1-S7

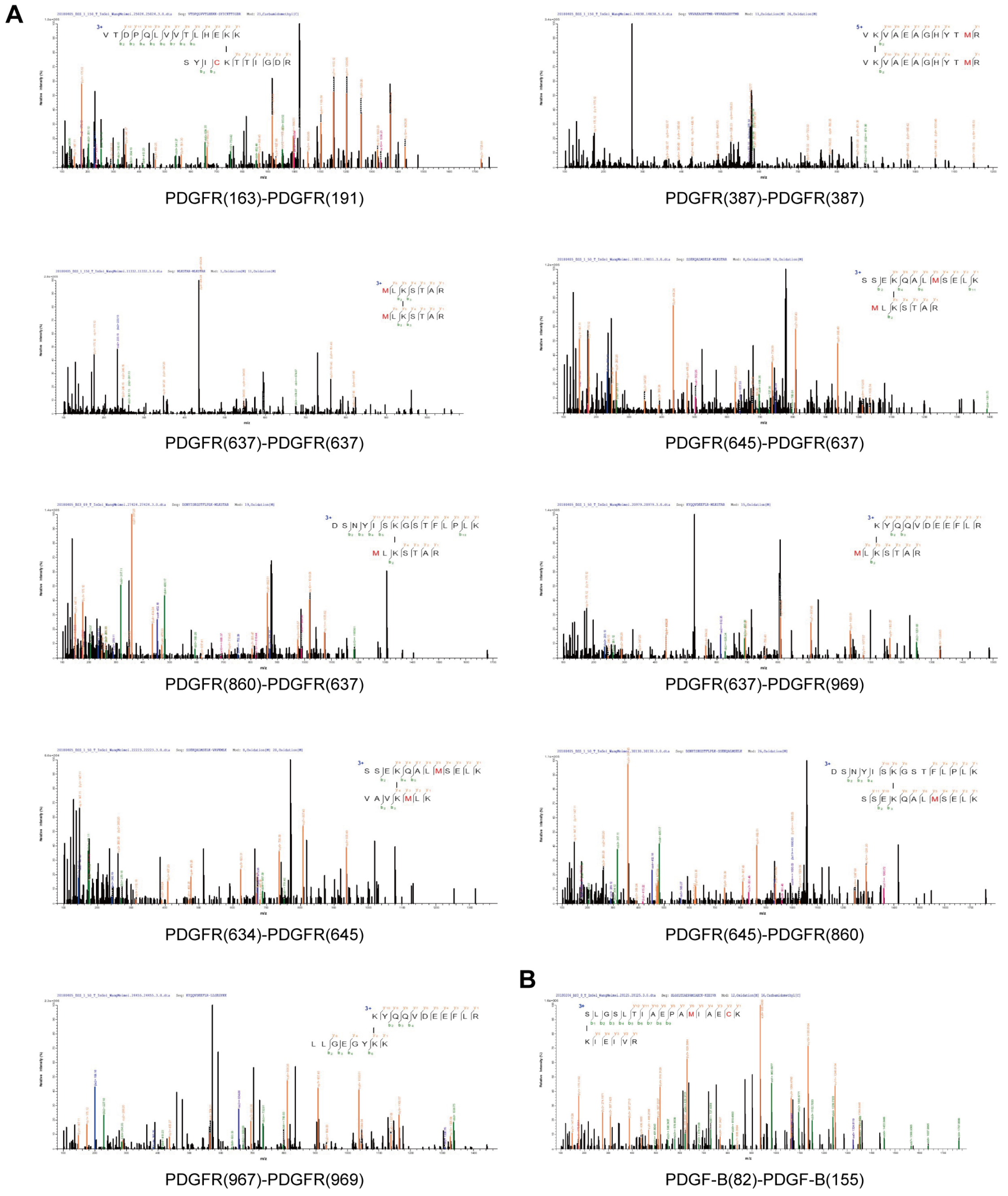


**Figure S1. Cross-linking spectra of activated PDGFRβ kinase dimer.** Purified PDGFRβ was treated with PDGF-BB and cross-linked with $BS^3$. (A) Cross-linked peptides of PDGF-BB-bound PDGFRβ. (B) Cross-linked peptide of PDGF-BB.

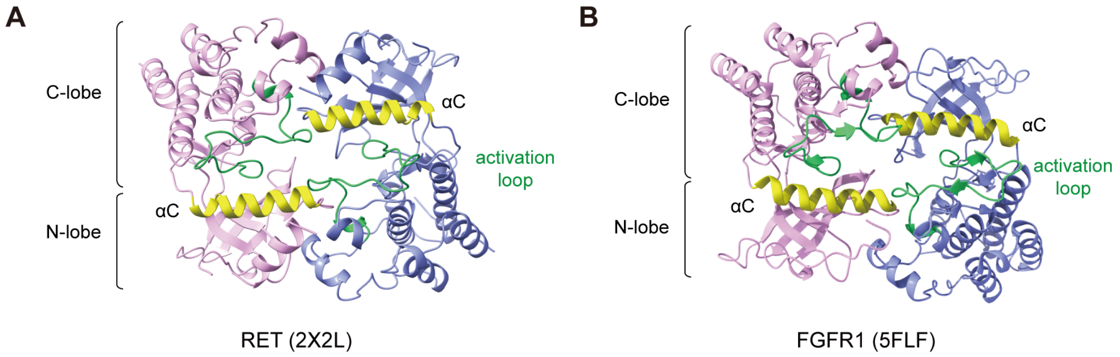


**Figure S2. Similar symmetric assembly of kinase dimer.** The symmetric dimer assembly was conserved among the type III receptor tyrosine kinases. (A) The crystal packing of the activated RET kinases (PDB 2X2L). (B) The crystal packing of activated FGFR1 kinases (PDB 5FLF).

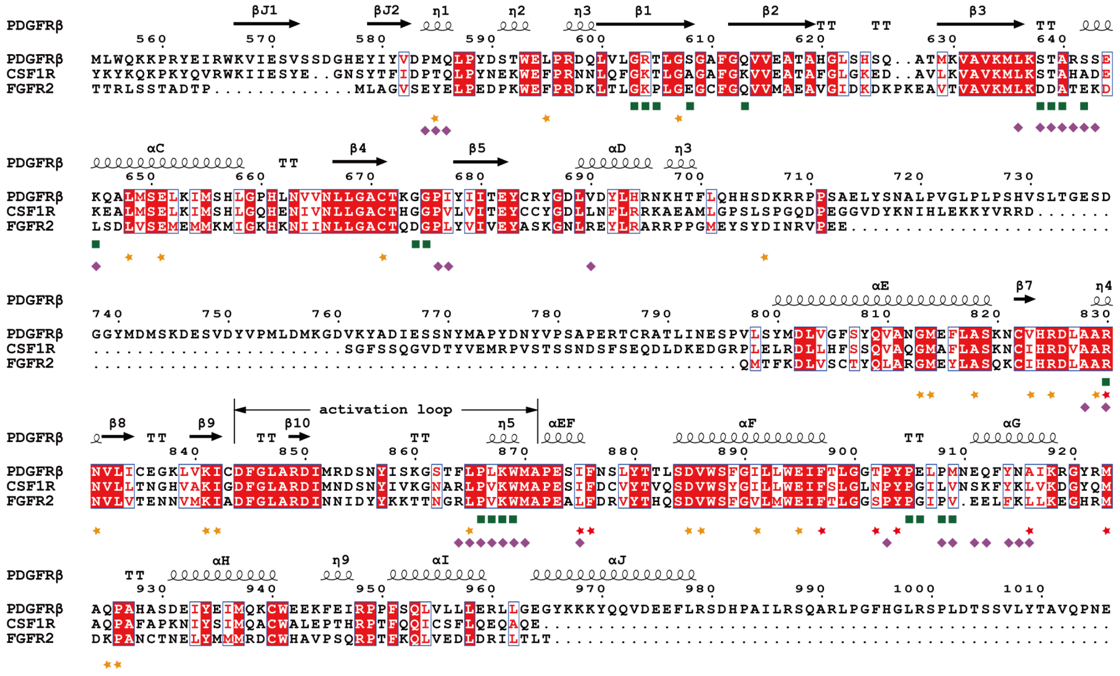

**Figure S3. Sequence alignment of PDGFRβ, CSF1R and FGFR2**. Residues at the kinase dimerization interface of PDGFRβ and FGFR2 were marked with green squares and purple diamonds, respectively. Residues at CSF1R kinase dimerization interface were marked with yellow and red stars with red stars emphasizing the mutations associated with HDLS.

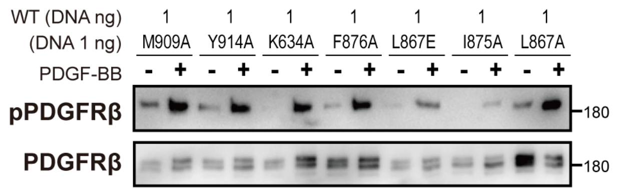


**Figure S4. Kinase activities of co-transfected WT PDGFRβ with different mutants.** In the transfection, the amounts of the plasmid encoding WT PDGFRβ and the plasmid encoding the mutant were kept at equal mass ratio. The phosphorylation and protein expression levels were detected by western blotting using 4G10 and protein C antibodies, respectively.

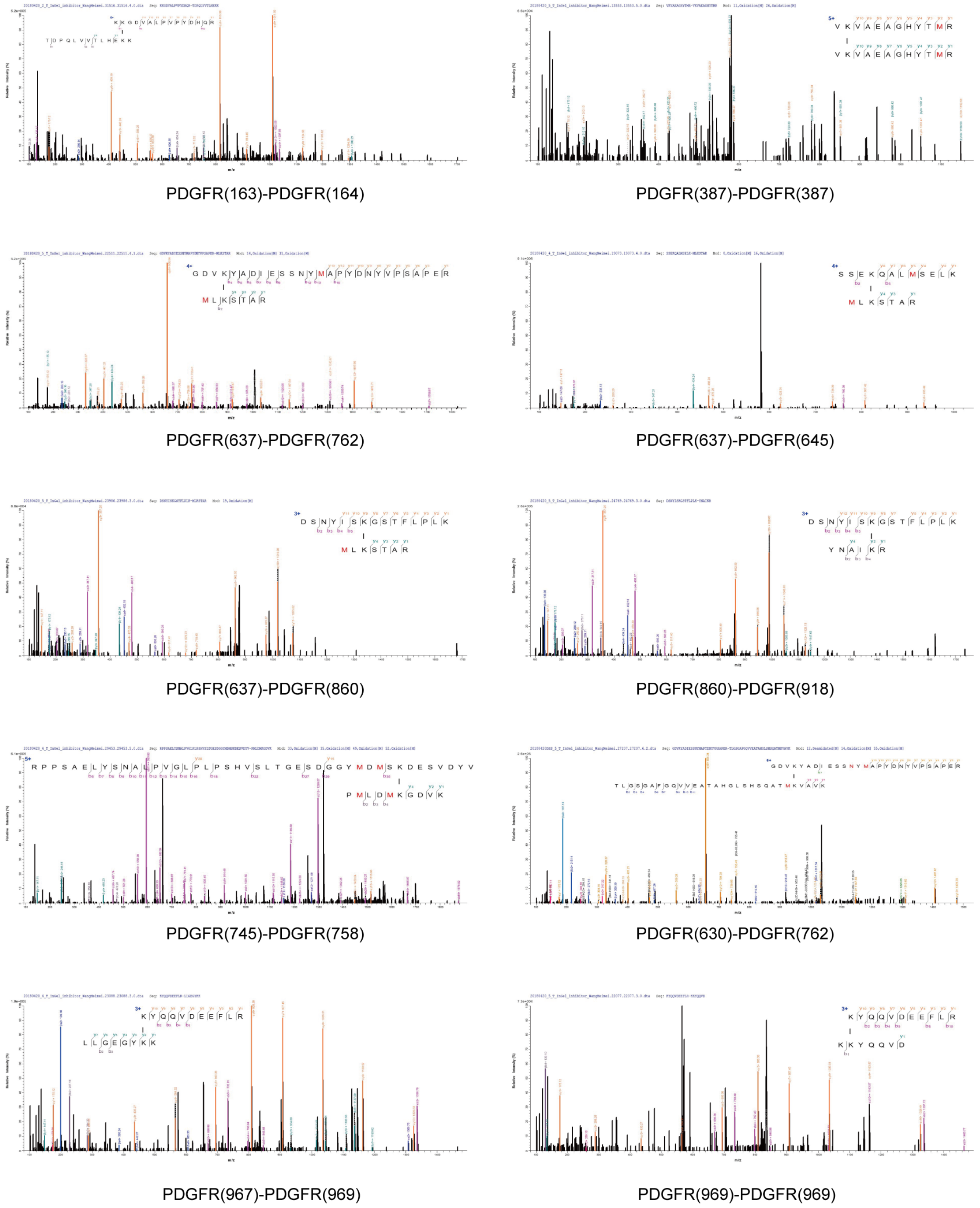


**Figure S5. Cross-linking spectra of inactive kinase dimer of PDGFRβ.** Purified PDGFRβ was treated with PDGF-BB and Dovitinib to stabilize the kinases in ligand-bound receptors in an inactive conformation. Then, the sample was cross-linked with $BS^3$.

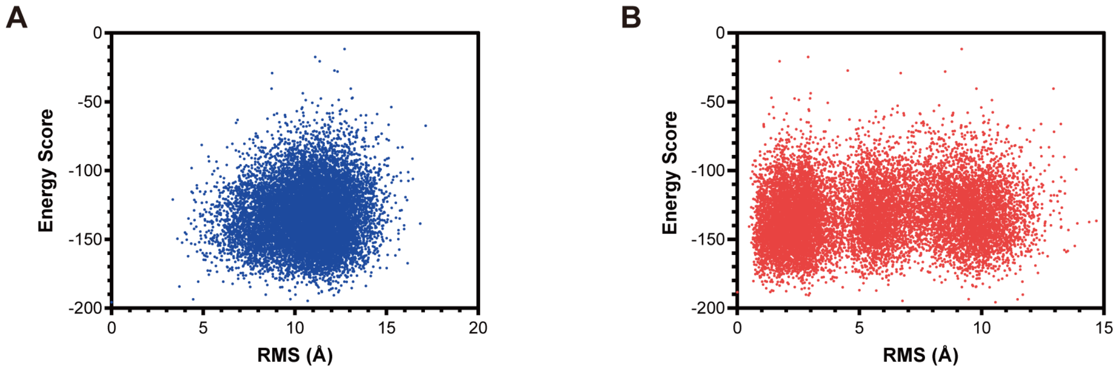


**Figure S6. Energy Score *vs* RMS in the *Ab initio* modeling of PDGFRβ kinase insert using Rosetta.** All-atom models of PDGFRβ kinase insert were generated and refined using Rosetta. (A) The plot of energy score *vs* RMS for PDGFRβ kinase insert models (B) The plot of energy score *vs* RMS for PDGFRβ kinase insert core structures. The RMSs for the top cluster of kinase insert models were converged to 8 Å, while the core structures were converged to 1 Å.

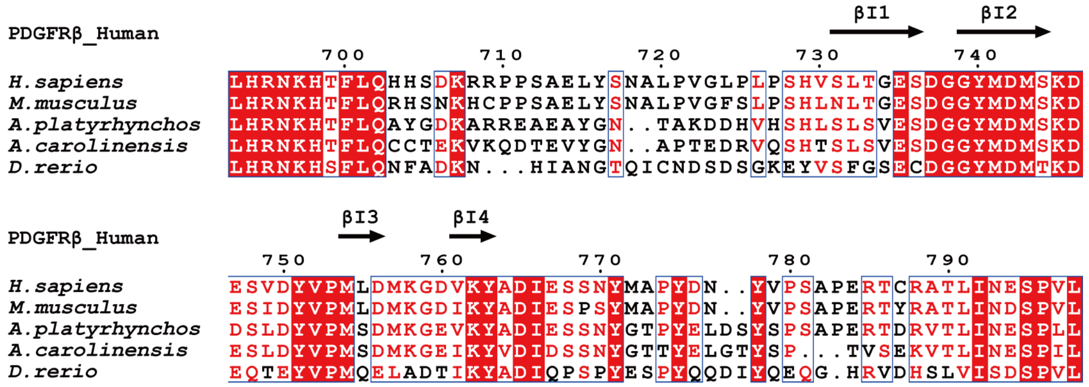

**Figure S7. Sequence alignment of PDGFRβ kinase insert core.** The sequence of PDGFRβ kinase insert core is conserved across Metazoa.
